# Diffusion Basis Spectrum Imaging with Deep Neural Network Differentiates Distinct Histology in Pediatric Brain Tumors

**DOI:** 10.1101/2020.04.02.020875

**Authors:** Zezhong Ye, Komal Srinivasa, Joshua Lin, Jeffrey D. Viox, Chunyu Song, Anthony T. Wu, Peng Sun, Sheng-Kwei Song, Sonika Dahiya, Joshua B. Rubin

## Abstract

High-grade pediatric brain tumors constitute the highest mortality of cancer-death in children. While conventional MRI has been widely adopted for examining pediatric high-grade brain tumor clinically, accurate neuroimaging detection and differentiation of tumor histopathology for improved diagnosis, surgical planning, and treatment evaluation, remains an unmet need in the clinical management of pediatric brain tumor. We employed a novel Diffusion Histology Imaging (DHI) approach that incorporates diffusion basis spectrum imaging (DBSI) and deep neural network. DHI aims to detect, differentiate, and quantify heterogenous areas in pediatric high-grade brain tumors, which include normal white matter (WM), densely cellular tumor (DC tumor), less densely cellular tumor (LDC tumor), infiltrating edge, necrosis, and hemorrhage. Distinct diffusion metric combination would thus indicate the unique distributions of each distinct tumor histology features. DHI, by incorporating DBSI metrics and the deep neural network algorithm, classified pediatric tumor histology with an overall accuracy of 83.3%. Receiver operating analysis (ROC) analysis suggested DHI’s great capability in distinguishing individual tumor histology with AUC values (95%CI) of 0.983 (0.985-0.989), 0.961 (0.957-0.964), 0.993 (0.992-0.994), 0.953 (0.947-0.958), 0.974 (0.970-0.978) and 0.980 (0.977-0.983) for normal WM, DC tumor, LDC tumor, infiltrating edge, necrosis and hemorrhage, respectively. Our results suggest that DBSI-DNN, or DHI, accurately characterized and classified multiple tumor histologic features in pediatric high-grade brain tumors. If further validated in patients, the novel DHI might emerge as a favorable alternative to the current neuroimaging techniques to better guide biopsy and resection as well as monitor therapeutic response in patients with high-grade brain tumors.

## Introduction

Pediatric brain tumors are the second most common childhood malignancy and the most common solid tumor in children (1). Pediatric brain cancer has unfortunately surpassed leukemia to become the most common cause of death from cancer in children in the US (2). It is estimated that 2,940 new cases of childhood (0-14 age group) and adolescent (15-19 age group) primary malignant and non-malignant central nervous system (CNS) tumors will be diagnosed in the United States (US) in 2020 (3).

In the past two decades, technological advances in neuroimaging have enabled clinicians to make earlier diagnosis to spot tumor recurrence or dissemination with more certainty (5). Yet these discoveries were tested on adult brain tumor patients; it is worthy to note that pediatric brain tumors do not exist on the same continuum as adult tumors. Rather, pediatric brain tumor represents a distinct group of tumors with unique genomic and imaging characteristics (6). Evaluating pediatric brain tumors is often a diagnostic challenge due to their diverse tumor pathologies, nonspecific or overlapping imaging findings, susceptibility artifacts from intratumoral calcification or hemorrhage, and motion artifacts in young children (7). Conventional MRI-based diagnoses also fail to offer adequate information regarding the specific tumor type, tumor grade, tumor viability, and treatment response of lesions. Although advanced MRI techniques like diffusion-weighted imaging (DWI), diffusion tensor imaging (DTI), perfusion MRI, MR spectroscopy (MRS), and susceptibility-weighted imaging (SWI), are incorporated into clinical MRI protocols, they still fall short (7–9). The multiparametric imaging (mpMRI) approach still often fail to accurately reflect tumor histopathology such as lesion cellular density, necrosis, hemorrhage, and infiltrative edges. As prior studies documented the histological and radiological tumor heterogeneity that exist within high grade tumor lesions, it is imperative to develop a technique capable of discerning the varied appearance of these lesions non-invasively(10).

We developed diffusion basis spectrum imaging (DBSI) (11) and demonstrated its ability to quantitatively characterize pathologies in multiple central nervous system diseases, including glioblastoma (12), multiple sclerosis (13–15), spinal cord injury (16), and epilepsy (17). We modified DBSI to better characterize these pathology-directed structural changes to ultimately incorporate DBSI-derived diffusion metrics with a deep neural network (DNN) algorithm to detect and differentiate various tumor histologic components in pediatric high-grade brain tumors. Here we introduce the novel diffusion histology imaging (DHI) technique.

## Results

### Patient characteristics

The 9 patients included in this study ranged from 4 to 17 years of age at the time of initial diagnosis. The mean age was 10.8 ± 3.7 years old. The patients’ age at autopsy ranged from 7 to 18, with a mean of 13.1 ± 3.7 years. Four tumors were located in the brainstem, two in the thalami, one in the right cerebral cortex, and one at the cerebellopontine angle. These were confirmed to be diffuse midline gliomas with H3K27M mutation by immunohistochemistry (n=4), glioblastoma (n=3), and embryonal tumor with multilayered rosettes with LIN28A protein overexpression (medulloepithelioma phenotype, NEC; n=1). Note one patient with neurofibromatosis 1 (NF1) had developed three different tumors at three distinct time points; these were an optic pathway glioma (pilocytic astrocytoma), a diffuse astrocytoma, WHO grade II involving the right parieto-temporal lobe and a CNS embryonal tumor involving the right temporal lobe. All details are summarized in Table 1.

**Table 1.** Patient information.

| Patient ID | Age at Diagnosis | Age at Post-mortem | Gender | Location | Histologic Diagnosis | Molecular Alterations |
| --- | --- | --- | --- | --- | --- | --- |
| WU-1 | 9 | 9 | F | Thalamus | Diffuse midline glioma, WHO grade IV | H3K 27M mutant (by immunohistochemistry) |
| WU-2 | 11 | 14 | M | Left temporal lobe | Glioblastoma, IDH wildtype, WHO grade IV | Tumor progressed from IDH wildtype anaplastic astrocytoma. Next generation sequencing showed *CREBBP* G1479 alteration |
| WU-3 | 11 | 12 | F | Right parietal-occipital lobe | Diffuse midline glioma, WHO grade IV | H3K 27M (by immunohistochemistry) |
| WU-4 | 7 | 16 | M | 1. Right temporal lobe | CNS embryonal tumor with anaplastic features, WHO grade IV (2013) | Background of NF1 with three tumors at different time points. 18 non-synonymous variants were identified by next generation sequencing *TP53*, p.R213Dfs*34, *TP53* and p.T211I *MAP2K2*,p.I369V |
|  |  |  |  | 2. Right posterior temporoparietal lobe | Diffuse astrocytoma, WHO grade II (2006) |  |
|  |  |  |  | 3. Optic pathway | Pilocytic astrocytoma, WHO grade I (not sampled until post-mortem) |  |
| WU-5 | 10 | 10 | M | Pons | Diffuse midline glioma, WHO grade IV | H3K 27M mutant (by immunohistochemistry) |
| WU-6 | 13 | 14 | M | Pons | Diffuse midline glioma, WHO grade IV | H3K 27M mutant (by immunohistochemistry) |
| WU-7 | 4 | 7 | F | Right cerebellopontine angle | Embryonal tumor with multilayered rosettes, medulloepithelioma phenotype, WHO grade IV, NEC | FISH could not demonstrate C19MC alteration but multifocal LIN28A protein expression was seen by immunohistochemistry |
| WU-8 | 17 | 18 | M | Right cerebral hemisphere (extensive involvement left side, brainstem and cerebellum) | Glioblastoma, IDH wildtype, WHO grade IV | Loss of 10q (*PTEN*) and monosomy 10 (by FISH); no *EGFR* amplification or polysomy of chromosome 7 |
| WU-9 | 15 | 18 | F | Epicenter in brainstem, right thalamus, right basal ganglia, and cerebellum | Glioblastoma, IDH wildtype, WHO grade IV | H3K 27M negative (by immunohistochemistry) |

### DBSI diffusion metric maps revealed tumor histology

Figures 1 and 2 show a representative case from a 16-year-old brain tumor patient with embryonal neoplasm (WHO Grade IV). Clinical gadolinium (Gd)-enhanced T1-weighted imaging of this patient several weeks prior to death, indicated a new lesion in the right temporal lobe of the brain (Fig. 1A, square). At autopsy, the brain was removed and immediately suspended in formalin for fixation (Fig. 1B). Coronal slices revealed a large hemorrhagic and necrotic tumor mass with its epicenter in the right thalamus (Fig. 1C). Tissue blocks were obtained from this region (Fig. 1D) for *ex vivo* imaging (Fig. 2). Of note, this patient had two other known tumors, one in his optic pathway (WHO grade I) and another diffuse astrocytoma (WHO grade II) in right posterior temporo-parietal lobe. The boundaries of latter were however relatively indistinct from the high-grade hemorrhagic and necrotic embryonal neoplasm (WHO grade IV) by gross examination alone.

**Figure 1.**
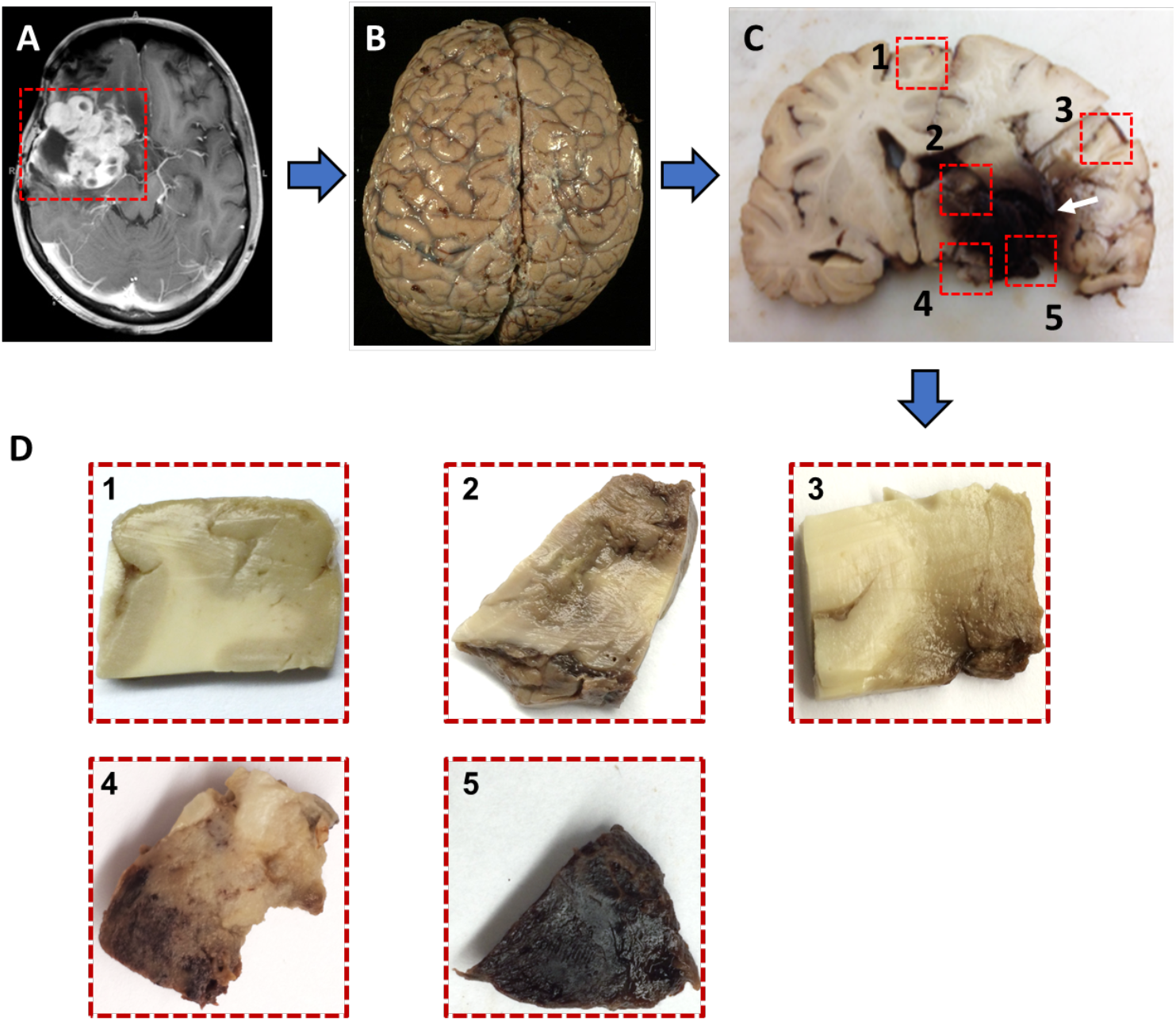
Illustration of brain specimen procurement from a patient with high-grade pediatric brain tumor. (**A**) *In vivo* Gd-enhanced T1-weighted image indicated a large lesion (square) with heterogeneous intensities in the right posterior region from a 16-year-old patient with embryonal neoplasm (WHO Grade IV). (**B**) Brain specimen was procured and immediately formalin-fixed. (**C**) Coronal slices revealed a large tumor with admixed hemorrhage and necrosis in the right thalamus (arrow). (**D**) Five tissue blocks were prepared in total i.e. from tumor (2, 4), tumor interface with normal adjacent brain (3), hemorrhage and necrosis (5), as well as grossly normal brain (1) (**C**).

**Figure 2.**
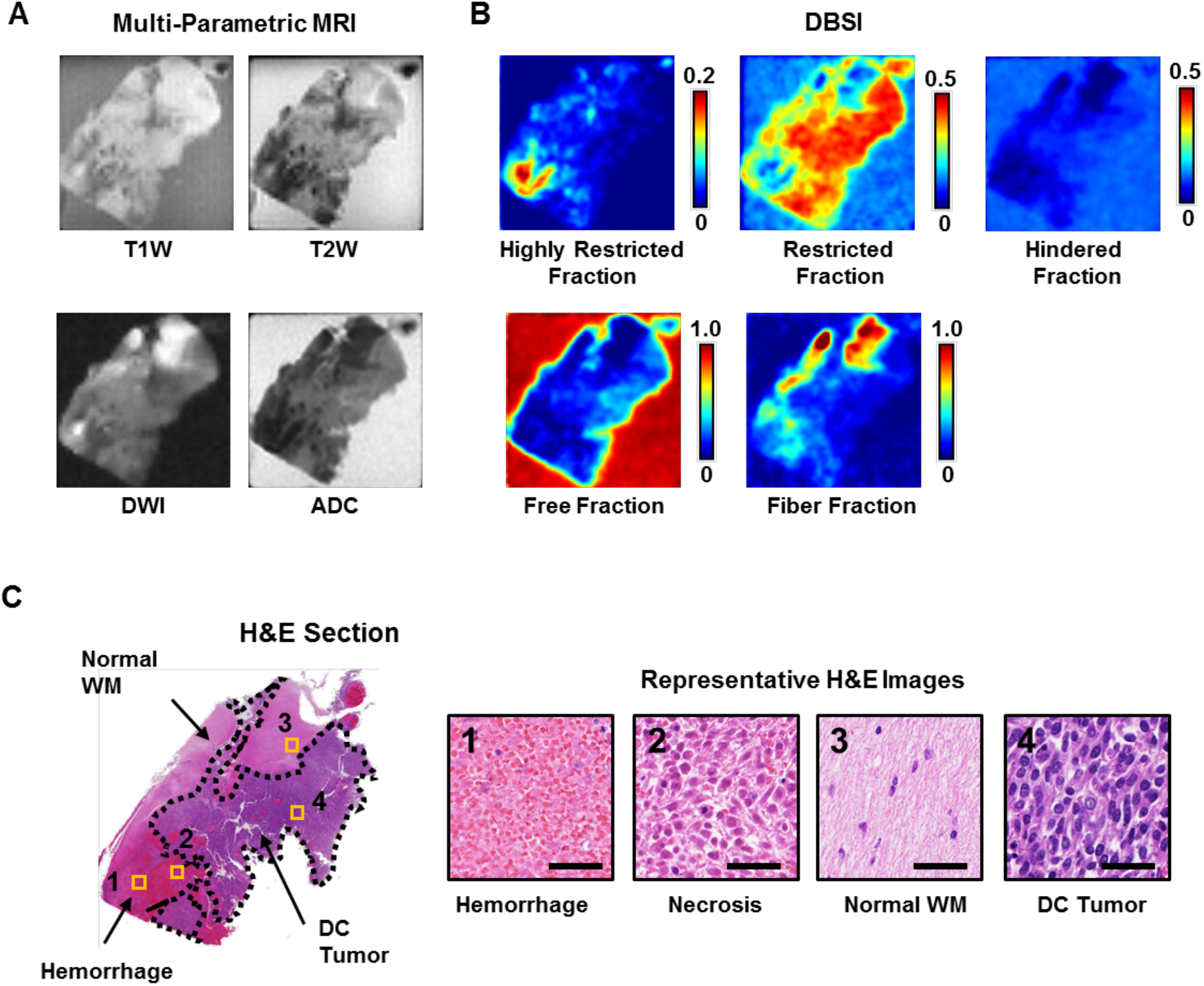
One tissue block was imaged with ex vivo MRI, followed by histologic processing and evaluation. **(A)** By referring to H&E image, densely cellular (DC) tumor region showed signal iso-intensity compared to normal white matter (WM) region in both T1W and T2W images. Hemorrhagic area showed signal hypointensity compared to other areas in both T1W and T2W images. Normal WM and hemorrhagic region showed hyperintensity in DWI and lower ADC values in ADC map. (**B**) Representative DBSI maps are shown for the same tissue block. Hyperintense areas in highly restricted fraction, restricted fraction and fiber fraction maps correlated with hemorrhage, densely cellular tumor and normal WM regions from corresponding H&E images. (**C**) H&E images demonstrating DC tumor, hemorrhage, necrosis and normal WM. Representative enlarged images from these regions are shown. WM, white matter. DC tumor, densely cellular tumor. Scale bar measures 50 um.

A densely cellular (DC) tumor region and normal white matter (WM) were indistinguishable in both T1WI and T2WI (Fig. 2A). Hemorrhage showed signal hypointensity compared to other regions in T1WI and T2WI (Fig. 2A). Normal WM and the hemorrhagic region showed higher DWI signal intensities and lower ADC values than DC tumor regions. This was in contrast to the clinical imaging in which the densely cellular tumor region was correlated with decreased ADC values. Representative DBSI diffusion maps are shown for the same tissue block (Fig. 2B). Regions with signal hyperintensity in highly restricted fraction, restricted fraction and fiber fraction maps correlated with hemorrhage, DC tumor and normal WM regions in the corresponding H&E images (Fig. 2C).

### Group analysis on diffusion metrics on different tumor histologic components

MR images were co-registered with H&E images on a voxel-to-voxel basis; imaging-voxels from segmentations of the five different tumor histologic components were subsequently obtained and plotted for group comparison (Fig. 3). Compared with normal white matter, DC tumor, LDC tumor, infiltrative edge, necrosis and hemorrhage show increased ADC values (Fig. 3A). The ADC of DC tumor (0.44 ± 0.19 μm^2^/ms) was higher than the infiltrating edge (0.29 ± 0.16 μm^2^/ms) but lower than LDC tumor (0.50 ± 0.26 μm^2^/ms) or necrosis (0.65 ± 0.37 μm^2^/ms). DTI fractional anisotropy (FA) values of tumor infiltration regions were similar s to normal white matter (0.24 ± 0.14 vs. 0.24 ± 0.11); both tumor infiltration and normal white matter FA values were substantially higher than that of the tumor histologic features (Fig. 3B). The isotropic ADC eliminated the signal contributions from anisotropic components such as white matter tracts. The comparison of isotropic ADC values among the distinct histologic features displayed similar relationship with mean ADC and were consistently higher than mean ADC (Fig. 3C). For the highly restricted fraction, normal WM showed much higher values than other histologic features. DC tumor and LDC tumor exhibited the lowest highly restricted fraction values among all histologic features (Fig. 3D). For the restricted fraction, DC tumor (0.34 ± 0.11) showed higher values than normal WM (0.27 ± 0.11), LDC tumor (0.29 ± 0.09), infiltrative edges (0.27 ± 0.11) and necrosis (0.23 ± 0.15). This result correlated well with the expected cellularity decrease from DC, LDC tumor regions to necrotic tissue (Fig. 3E). As expected, necrosis was characterized by higher hindered fraction (0.40 ± 0.22) and free fraction values (0.10 ± 0.12) than any of the other histologic features. In the anisotropic fraction, normal WM (0.37 ± 0.12) and infiltrative edge (0.37 ± 0.17) had similar values; this anisotropic component were much higher than other histologic components.

**Figure 3.**
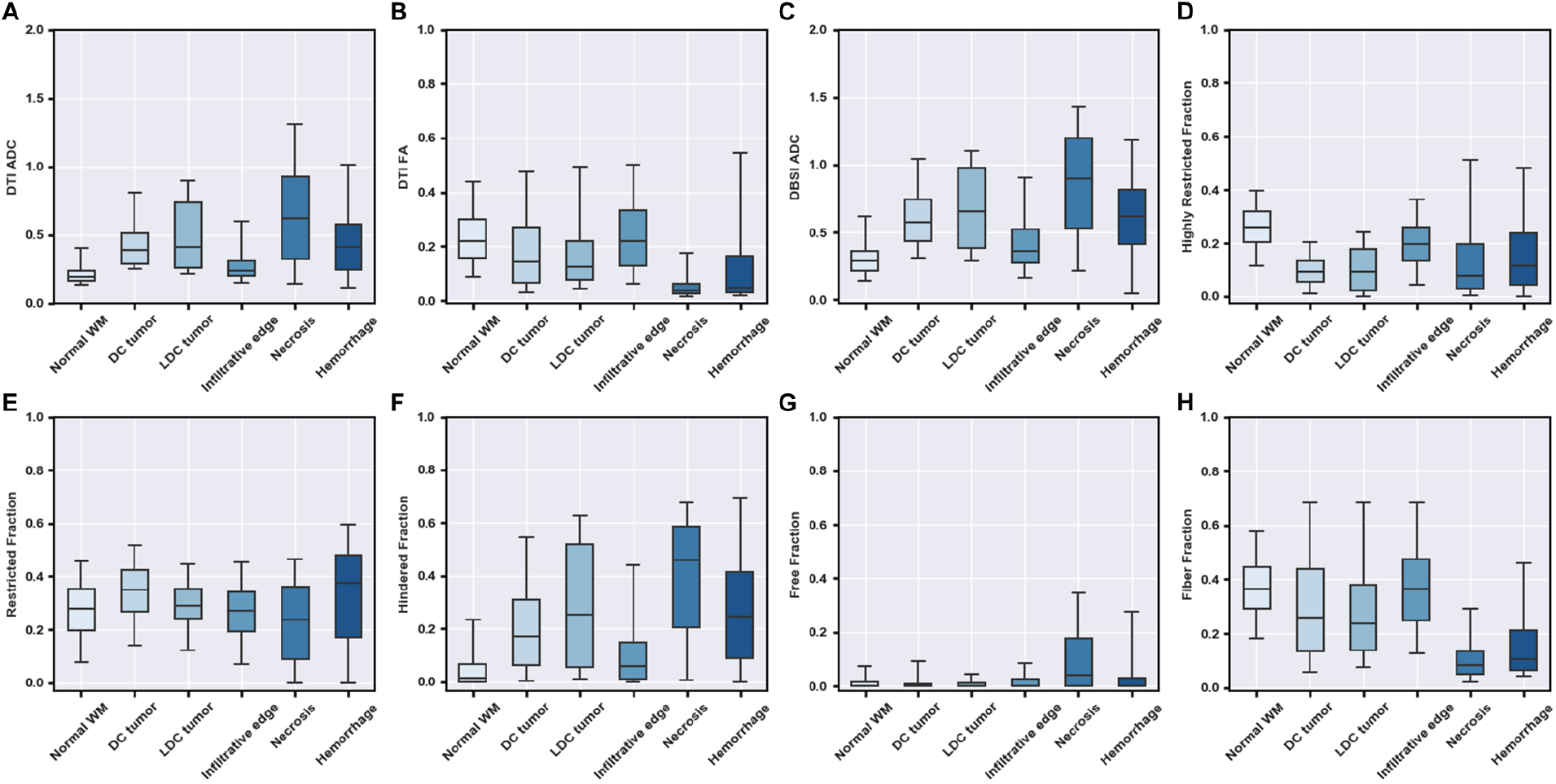
Group analysis on different tumor histologic components on representative diffusion metrics including (A) ADC, (B) DTI FA, (C) DBSI isotropic ADC, (D) highly restricted fraction, (E) restricted fraction, (F) hindered fraction, (G) free fraction, and (H) fiber fraction. Particularly, normal WM and the infiltrative edge showed higher fiber fraction and DTI-FA than the other tumor histologies. DC tumor and LDC tumor showed higher restricted fraction values than other histologies. Necrosis showed higher ADC, hindered fraction and free fraction values as well as lower restricted fraction, fiber fraction and DTI-FA compared to the other histologies. These findings were collectively consistent with DBSI’s modelling for malignant brain tumor. ADC, μm^2^/ms.

### Classifications of tumor histologic components

A total of 114,786 imaging voxels were used to train the DHI model after data balancing. We first performed a multi-class classification on the normal WM, DC tumor, LDC tumor, infiltrative edge, necrosis and hemorrhage regions. Representative H&E images with one MRI voxel size indicated distinct histologic features (Fig 4A). For the independent test set (n = 9,446), we achieved an overall accuracy of 83.3%. Confusion matrix analysis indicated strong concordance between DHI predictions and the neuropathologist-identified histologic features (Fig. 4B). DHI accurately predicted normal WM, DC tumor, LDC tumor, infiltrative edge, necrosis, and hemorrhage, with true positive rates of 86.4%, 81.5%, 95.2%, 75.2%, 78.1% and 83.9%, respectively.

**Figure 4.**
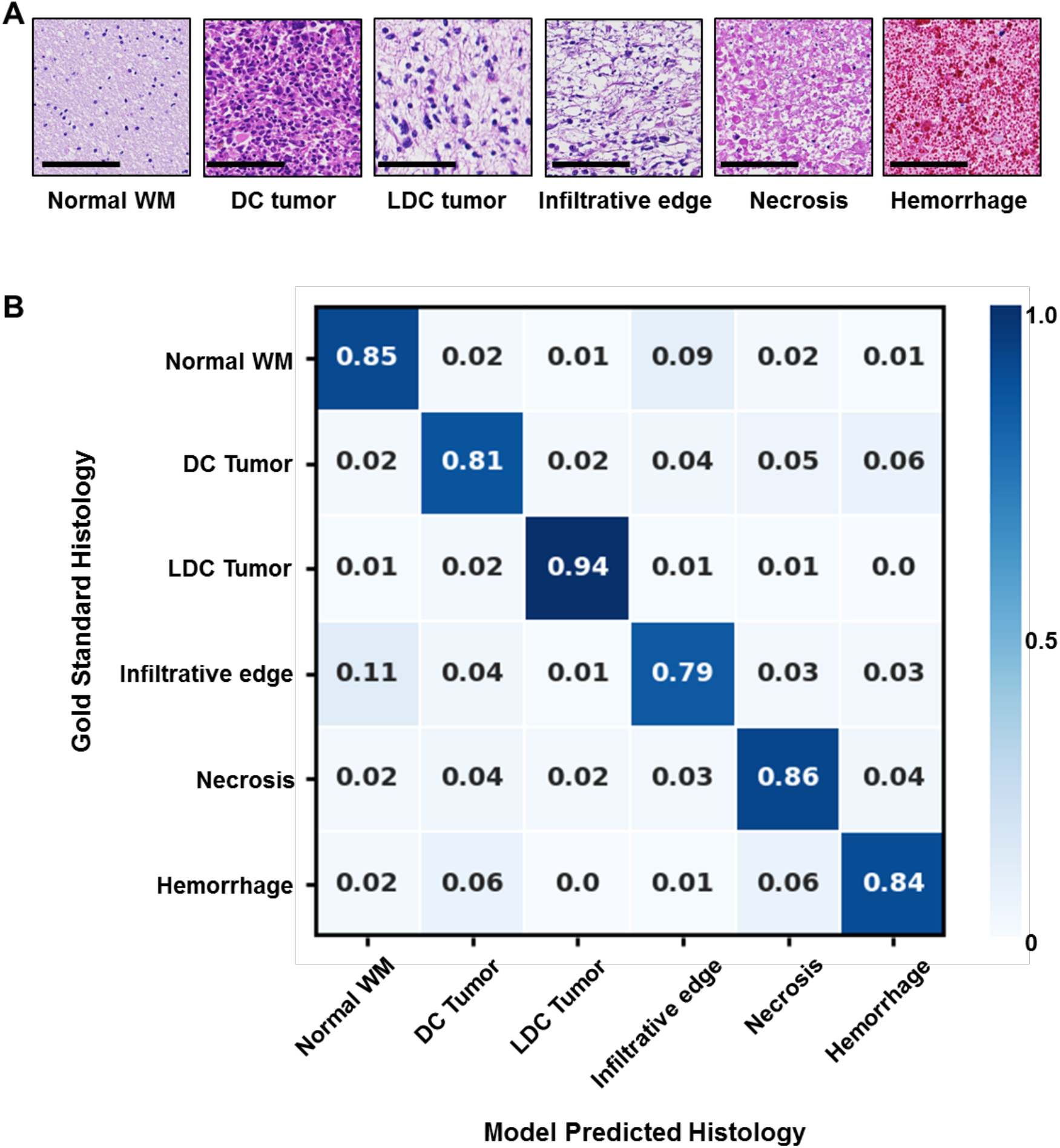
(**A**) Representative H&E images of normal white matter, densely cellular tumor, less densely cellular tumor, infiltrative edge, necrosis and hemorrhage, respectively. **(B)** Independent test dataset confusion matrix for the predictions of DHI versus gold standard, i.e. histologic examination (n=9,446). Rows contain tumor histologic classifications identified by a neuropathologist, and columns represent tumor histologic classifications as predicted by DHI. Scale bar measures 100 μm.

To test DHI’s ability to distinguish each individual tumor histology, we adopted a one-versus-rest strategy to perform ROC and precision-recall analysis (Fig. 5). The ROC curves indicated great AUC values in the differentiation of all six different histologic components. We calculated 95% confidence intervals (CI) of AUCs using the percentile bootstrap method with 10,000 iterations. The AUC values (95% CI) were 0.983 (0.985 – 0.989), 0.961 (0.957 – 0.964), 0.993 (0.992 – 0.994), 0.953 (0.947 – 0.958), 0.974 (0.970 – 0.978) and 0.980 (0.977 – 0.983) for normal WM, DC tumor, LDC tumor, infiltrative edge, necrosis and hemorrhage, respectively (Table 2). We also calculated sensitivity and specificity for each class under the Youden Index. All the sensitivity values were higher than 89% with specificity values higher than 87% (Table 2). We also calculated precision-recall (PR) curves and F_1_-scores to provide complementary information to address ROC analyses’ insensitivity to class imbalance and the possible overestimation of model performance, The PR curves performance inferiorly on tumor infiltration (Fig. 4D, AUC 0.796) and necrosis (Fig. 4E, AUC 0.876)) when compared to other tumor histologic regions. Similarly, the F_1_-scores of the infiltrative edge (0.703) and necrosis (0.753) were worse those of normal white matter (0.863), DC tumor (0.834), LDC tumor (0.930), and hemorrhage (0.833) (Table 2).

**Table 2.**
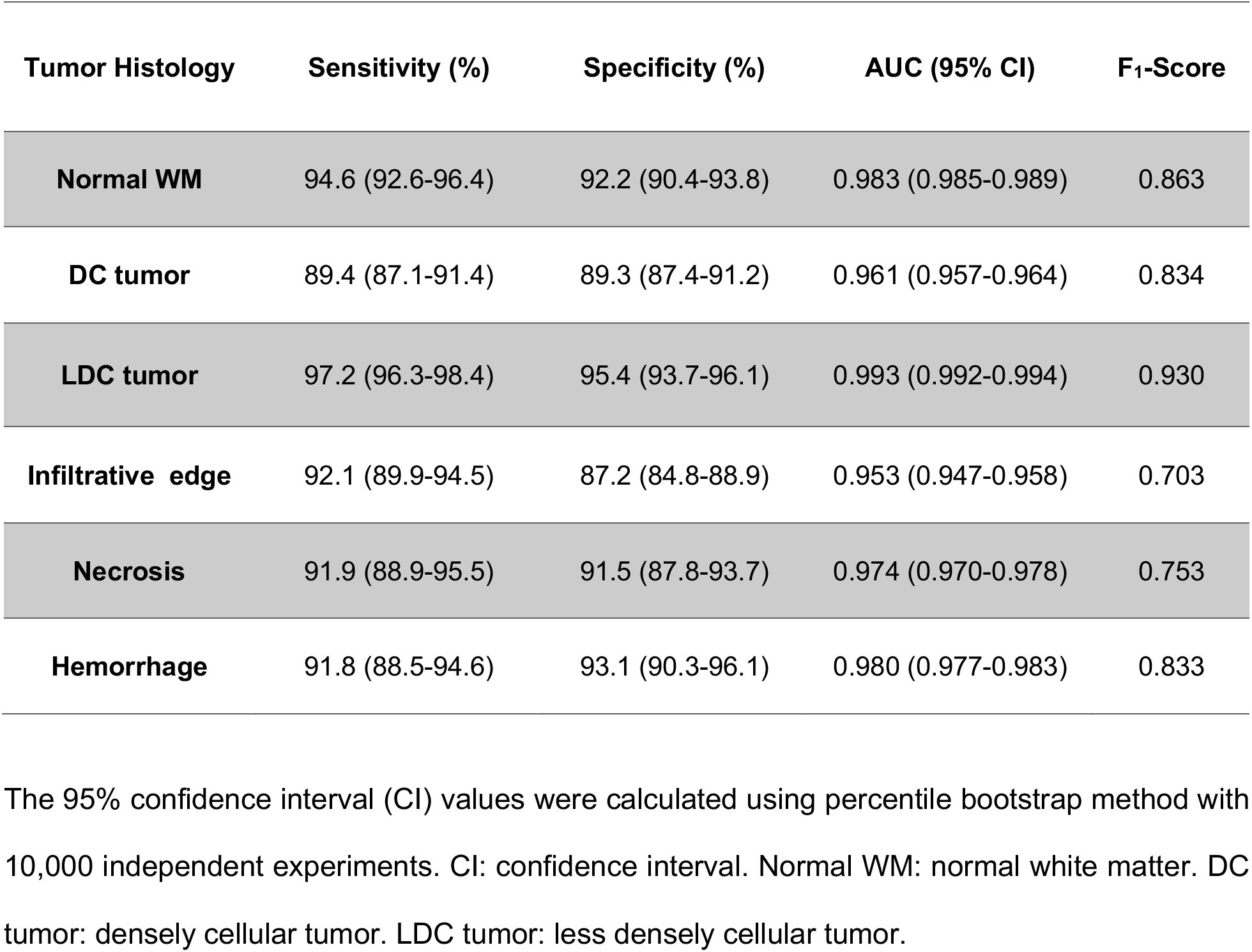
Diagnostic performances of DHI in classifying different tumor histologies.

**Figure 5.**
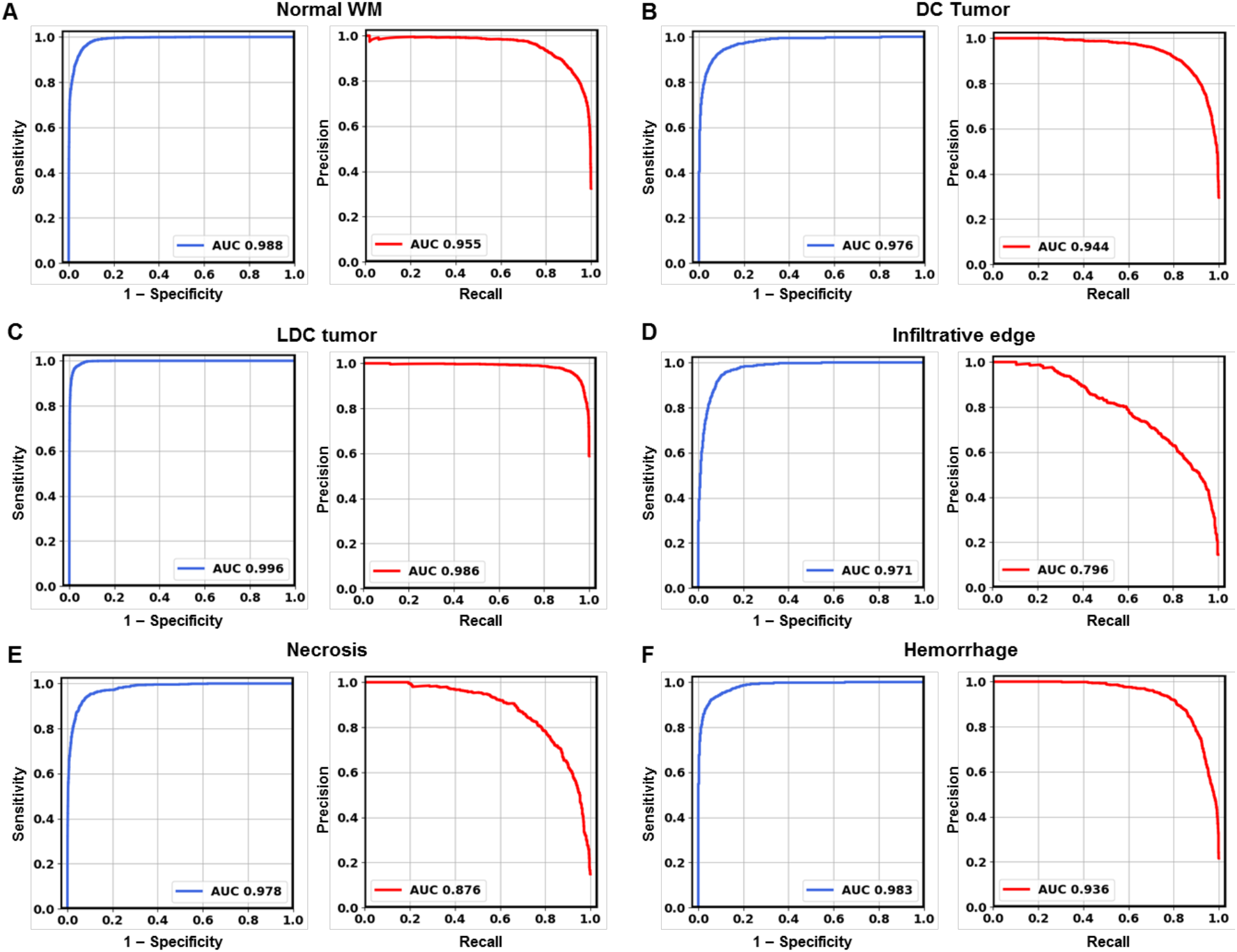
Receiver operating characteristics (ROC) curves and precision-recall (PR) curves calculated using one-vs-all strategy for 6 different tumor histological components including (A) normal white matter, (B) densely cellular tumor, (C) less densely cellular tumor, (D) tumor infiltrative edge, (E) tumor necrosis and (F) hemorrhage. All 6 ROC curves showed high areas under curve (AUC), indicating strong sensitivity and specificity in detecting these tumor histologic components. Tumor infiltrative edge did not perform as well as other histologic components in precision-recall analysis, indicating that tumor infiltration could be overestimated by the model. WM, white matter. DC tumor, densely cellular tumor. LDC tumor, less densely cellular tumor.

## Discussion

Pediatric brain tumors are the leading cause of cancer-related childhood death. Current curative approaches in management rely, in most cases on complete surgical resection, followed by irradiation therapy and chemotherapy (5, 18). Histologic assessment of tumor cellularity, infiltration and necrosis is critical in diagnosis and grading, as well as subsequent clinical decision-making for patient management and follow-up (19). The current clinical gold standard, i.e. histologic examination, requires stereotactic biopsy or surgical resection (20), which carries potential risks including infections, seizures, stroke, coma, as well as brain swelling or bleeding (21). Sometimes inconclusive pathological findings result from inadequate samples would require the patient to endure another procedure (22). The repeated biopsies may further induce intracranial hemorrhage, increase risk of inpatient mortality and hospital disposition (22). An noninvasive neuroimaging approach is thus necessary to facilitate diagnosis or guide surgical plane in order to ensure better treatment response assessment, ultimately improving patient care (23).

While MRI remains the most common clinical imaging technique for evaluating CNS tumors (6), conventional MRI sequences such as T1WI and T2WI correlated poorly with tumor pathology of high-grade brain tumors because of the complicated histologic heterogeneity. For example, hyperintense regions in T2W and FLAIR images surrounding the enhancing tumor core cannot distinguish between infiltrative tumor, vasogenic edema, or immune cell (24). Gd enhancement in T1WI also could occur due to either tumor progression or radiation necrosis (25). Furthermore, conventional T1WI and T2WI imaging contrasts vary from scan to scan and are not quantitative, as they depend not only on the MR characteristics of brain tissue, but also on the scanner models, magnet strength, and pulse sequences. The diagnostic power of conventional MRI continues to be hinder by its diagnostic accuracy due to the many acquisition variables that prevents a quantitative diagnostic standard from emerging.

To address the limitations of conventional MRI and bridge the gap between histology and MRI for pediatric brain tumor diagnoses, we developed a novel imaging technique – DBSI. DBSI provides a simple tensor expression value to visualize morphological features resulting from both tumor and non-tumor elements of the brain that are indistinguishable by conventional MRI. In our previous studies, we demonstrated how DBSI derived restricted fraction not only correctly delineated tumor distribution in GBM specimens, but also the restricted fraction’s strong positive correlation with tumor cellularity identified on H&E histology (12). In this study, we demonstrated that the hyperintense restricted fraction regions accurately identify densely cellular tumor areas (Fig. 2). Group analysis across multiple samples with various tumor types also indicated DC tumor had higher restricted fraction values than either normal WM, LDC tumor, infiltrative edges or necrosis. From the areas of necrosis, infiltrating edge, LDC tumor and DC tumor, we observed a trend of gradually increasing restricted fraction value across these four types of histological areas, respectively. This indicated that the restricted fraction could serve as an accurate biomarker to assess tumor cellularity in high-grade pediatric brain tumors. In addition, necrosis showed much higher values in hindered fraction and free fraction than the other histologic components, indicating that these two diffusion metrics could effectively assess tumor necrosis. Furthermore, our results showed comparable ADC, FA, restricted fraction, and fiber fraction values between the infiltrative edge and normal WM, suggesting that these diffusion metrics lack adequate sensitivity to detect cellularity or white matter changes. Note that infiltrative edge showed higher isotropic ADC and hindered fraction values than do normal WM, potentially pointing to how tumor infiltration displaces normal parenchyma (26), destructs of white matter tracts (27), and/or forming vasogenic edema to disrupt blood brain barrier (28).

The combination of multiple DBSI spectrums highlights the difference within tumor histology, as illustrated by the group analysis of DBSI. The incorporation of different diffusion metrics significantly improves the machine learning framework to better quantitatively monitor the morphological changes in different tumor histologies. In this study, we demonstrated that DBSI-DNN could differentiate 6 major types of tumor histologic components with an overall accuracy of 83.3%. In detecting and distinguishing individual tumor histology, ROC analysis of our model calculated the AUC, sensitivity and specificity values of all identified histologic area types to be higher than 0.950, 89.4% and 87.2%, respectively. In the precision-recall analysis, the prediction of infiltrative edge was relatively low for precision-recall AUC (0.796) and in F_1_-score (0.703), which were likely due to the highly variable degrees of infiltration or inherent cellularity of the different tumors. For example, infiltrative edges with mild to intermediate tumor cellularity could be falsely predicted to be normal WM. Similar phenomenon was observed from the results of confusion matrix (Fig. 4B).

In contrast to previous studies, we adopted a voxel-wise analysis through precise co-registration between histology and MR images, which provides a better approach to bridging MRI and histology. Application of this approach accurately detected regions of diverse pediatric brain tumor that were enriched for histological heterogeneity (29, 30). Histologically, distinct voxels taken from a region from a specimen could be very drastically different from each other. Such heterogeneity possess challenge on imaging, however, since DBSI models each image voxel independently (11), DBSI provides an unique opportunity to assess the heterogeneous histologic features of tumors. DBSI-derived structural metrics are thus ideal to serve as the unique histologic features for machine learning. To the best of our knowledge, such a voxel-level validation on imaging markers for pediatric brain tumor histology is the first of its kind. Patient-wise analysis has been typically studied by correlate image metrics with clinical scores or survival rates. There have been attempts to correlate MRI lesions with tumor histopathology using stereotactic biopsy (31, 32). However, the distinct spatial and volume differences between MRI lesions and biopsy combined with the lack of co-registration made the results less reliable given the high histological heterogeneity of high-grade pediatric brain tumor.

There were several limitations of this study. First, the relatively small number of subjects (n = 9) of our series limited the broad applicability of the results. However, we performed voxel-wise analyses of a total of 94,453 imaging voxels from 45 brain specimens containing different areas of the brain to address the limitation on sample size. We performed a voxel-based modelling and computation to derive DBSI metrics, which avoids the issues concerning heterogeneity of high-grade brain tumor. Secondly, the unbalanced data distribution amongst different tumor histologic components imposed another limit since the imbalance could compromise the performance of a DNN model. We addressed the concern by employing an oversampling approach to balance the training data and adopted precision-recall analyses to provide complement ROC analyses. Thirdly, since this study was based on data from a single institution using the same scanner, we will further investigate this DHI’s efficacy by examine classification models across different scanner platforms and acquisition parameter variations in the future.

In conclusion, we have demonstrated DHI, which incorporates DNN, to accurately characterize and classify multiple tumor histologic components in pediatric high-grade brain tumors. While precise prediction of infiltrative edges was suboptimal, the collective findings are encouraging once validated with a larger sample size. With further, and more extensive, validations, these findings would provide a rationale to prospectively test DHI’s aiding potential on targeted biopsies in different brain regions, offering a potentially improved tumor therapeutic response assessment, a more accurate diagnostic yields to optimize treatment decisions.

## Methods

### Patient information

Nine post-mortem pediatric brain tumor specimens that were part of the Washington University Legacy Project were included for the study. Among these 9 pediatric patients, four were male and five were female. The institutional review board of Washington University School of Medicine approved the study.

### Postmortem brain specimen

A total of 45 samples were resected from tumor, tumor interface with normal adjacent brain, areas of hemorrhage and necrosis, as well as normal brain tissue (Fig. 1). The average size of the specimens was 8 mm ± 4 mm.

### *Ex vivo* MRI of brain specimen

Brain tumor specimens were submersed in formalin for ex vivo imaging to keep tissue hydration during study. The specimens were examined using a 4.7-T Agilent/Varian MR scanner (Agilent Technologies, Santa Clara, CA) and a custom-built circular surface coil (3.5-cm diameter). A multi-echo spin-echo diffusion weighted sequence with 99 diffusion-encoding directions with maximum b-values=3000 s/mm^2^ was employed to acquire DW images. The imaging parameters were as follows: repetition time (TR)=1500 ms, echo time (TE)=40 ms, time between application of gradient pulse 20 ms, diffusion gradient on time 8 ms, slice thickness 0.5 mm, field-of-view 24×24 mm^2^, data matrix 96×96, number of average 1, in-plane resolution 0.25×0.25 mm^2^. T2W images were acquired with a multi-slice spin echo sequence with TR=4000 ms, echo time TE=80 ms, field of view field-of-view 2.4×2.4 mm^2^, data matrix 96×96. T1W images were acquired with a gradient echo sequence with TR=80 ms, TE=10 ms, field of view field-of-view 2.4×2.4 cm^2^, 8 averages, data matrix 96×96.

### DBSI analysis of brain tumor

DBSI models brain tumor diffusion-weighted MRI signals as a linear combination of discrete multiple anisotropic diffusion tensors and a spectrum of isotropic diffusion tensors:

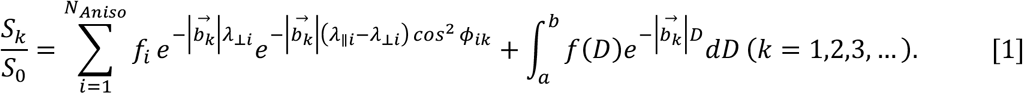

In [1], *b_k_* is the *k*^th^ diffusion gradient; *S_k_/S_0_* is the acquired diffusion-weighted signal at direction of *b_k_* normalized to non-diffusion-weighted signal; *N_Aniso_* is number of anisotropic tensors to be determined; *ϕ*_*ik*_ is the angle between diffusion gradient (*b_k_)* and principal direction of the *i^th^* anisotropic tensor; 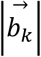 is *b*-value of the *k^th^* diffusion gradient; λ_*Ⅱi*_ and _*⊥i*_ are axial and radial diffusivity of the *i^th^* anisotropic tensor under the assumption of cylindrical symmetry; *f_i_* is signal-intensity-fraction of the *i^th^* anisotropic tensor; *a, b* are low and high diffusivity limits of isotropic diffusion spectrum; *f(D)* is signal-intensity-fraction at isotropic diffusivity *D*.

Based on our *ex vivo* MRI and histological analyses of resected specimens from previous studies (12), the following isotropic-diffusion profiles have been established based on diffusivity. We observed that highly restricted isotropic diffusion (0 ≤ D ≤ 0.2 μm^2^/ms) is associated with lymphocytes; restricted-isotropic diffusion (0.2 < D ≤ 0.8 μm^2^/ms) is associated with dense tumor cellularity; and hindered-isotropic diffusion (0.8 < D ≤ 2 μm^2^/ms) is associated with tumor necrosis. DBSI provides a simple tensor expression for individual image voxels to visualize morphological features secondary to tumor formation, some of which are not as discretely detectable by conventional MRI. It is the sensitivity of diffusion-weighted MRI signal to the microstructural changes that allows DBSI to more precisely reflect morphological changes resulting from tumor presence or other pathologic alterations. By using this feature of DBSI as the input for machine learning algorithms, we created DHI to recapitulate histopathologic analysis using MRI.

### Histologic staining and evaluation

The formalin-fixed tissue was embedded in paraffin after scanning. The paraffin embedded tissue was then sequentially sectioned at 5-μm thickness and stained with hematoxylin and eosin (H&E). Histology slides were digitized using NanoZoomer 2.0-HT System (Hamamatsu, Japan) with a 20× objectives for analyses. Two neuropathologists (KS and SD) reviewed all the histological slides with a consensus on the selected tumor histopathologic features. Regions of normal white matter (WM), densely cellular tumor (DC tumor), less densely cellular tumor (LDC tumor), necrosis, tumor infiltrative edge, and hemorrhage were outlined and drawn on H&E images with 20× magnification.

### Image processing

Voxel-wise DTI and DBSI analyses were performed by an in-house software developed using MATLAB^®^ (MathWorks; Natick, MA). To co-register H&E images and MR images, we first ensured that the plane of histology section of the brain tumor specimens matched closely with the slice plane of the MR images. We then performed a linear registration to co-register the images using an in-house software developed using MATLAB^®^. Pathologists defined regions were transferred to MR images using ITK-SNAP (http://www.itksnap.org/) (33).

### Deep neural network (DNN) model development and optimization

Our complete dataset consisted of 94,453 imaging voxels from 45 specimens obtained from 9 patients. The collected voxels were split into training, validation, and test datasets with a 8:1:1 ratio, respectively. Imaging voxels from test datasets were separated and distinct from the ones that were used in the training and validation steps. Validation set was employed to fine tune the model hyper-parameters. To balance data from groups of different tumor histologic components, a synthetic minority oversampling technique (SMOTE) (34) was applied to over-sample the minority group by introducing synthetic feature samples. This data balancing approach has been demonstrated to be beneficial for avoiding over-fitting and improving model generalization (34, 35). Data balancing were only applied to the training dataset, while the validation and test dataset was kept unchanged. The diffusion metrics assessed with our DNN modeling included 10 diffusion metrics provided from DBSI. Specifically, DBSI metrics include mean ADC, mean FA, fiber fraction, fiber fractional anisotropy (FA), fiber axial diffusivity (AD), fiber radial diffusivity (RD), restricted isotropic diffusion fraction (restricted fraction), restricted isotropic diffusivity, hindered isotropic diffusion fraction (hindered fraction), hindered isotropic diffusivity, free isotropic diffusion fraction (free fraction), free isotropic diffusivity.

A supervised deep neural network (DNN) was adopted to detect and classify tumor histologic components by referencing the H&E findings. The DNN model was developed using TensorFlow 2.0 framework in Python (36). In general, the DNN model was equipped with ten fully connected hidden layers. Batch normalization layer with a mini-batch size of 200 was used before feeding data to the next hidden layer to improve model optimization and prevent overfitting. Exponential linear units (37) were adopted to activate specific functions in each hidden layer. The final layer was a fully connected softmax layer that generated a likelihood distribution of six output classes. We used Adam optimizer with the default parameters of β_1_=0.9, β_2_=0.999 and mini-batch size of 200. The learning rate was manually tuned to achieve the fastest convergence. We chose cross-entropy as the loss function and trained the model to minimize the error rate on the validation dataset. Overall, hyper-parameters for the DNN architecture and optimization algorithm were chosen through a combination of grid search and manual tuning.

### Statistical analysis

Data represent mean ± SEM. In multi-class classification, confusion matrices were calculated and used to illustrate the specific examples of tumor histologic components where the model prediction agrees with the pathologists’ diagnoses. We also used one-versus-rest strategy to perform receiver operating characteristics (ROC) analysis. Area under curve (AUC) was calculated to assess model discrimination of each tumor histological component. Sensitivity and specificity values were calculated using Youden Index (38). The precision-recall curve and F_1_-scores were also calculated to provides complementary information to the ROC curves. F_1_-score (ranges from 0 to 1) favors models that maximize both precision and recall simultaneously, which is especially helpful to address the insensitivity of AUC on class imbalance. The 95% confidence interval values were calculated using the percentile bootstrap method with 10,000 independent experiments (39). All the statistical metrics and curves were calculated using the packages from Scikit-learn (40).

## Acknowledgements

This work was supported by The Taylor Rozier Hope for a Cure Foundation (JBR), The Josie Foundation (JBR), Matt’s Hats Foundation (JBR), The Derek Griffitts Foundation (JBR), and The Kewsi Prince Foundation (JBR), and in part by NIH R01-NS047592, P01-NS059560, U01-EY025500, and Department of Defense Idea Award W81XWH-12-1-0457. The authors are indebted to the patients and their families for donating the brains without which this research would not have been possible.

## Author contributions

ZY, KS and JL wrote the manuscript. ZY, KS and JV performed the experiments. ZY, JL, PS, and CS analyzed the experimental data. SD and KS performed the histopathologic evaluations. ZY and ATW developed the classifier and conducted the statistical analyses. JBR and SD provided surgical specimens for imaging. SD, JBR, and SKS interpreted data, designed and supervised the research, and supervised the writing of the manuscript. All authors have reviewed and approved the final version of the manuscript.

## References

1. Kline NE, and Sevier N. Solid tumors in children. Journal of Pediatric Nursing. 2003;18(2):96–102.

2. Curtin SC, Minino AM, and Anderson RN. Declines in Cancer Death Rates Among Children and Adolescents in the United States, 1999-2014. NCHS Data Brief. 2016(257):1–8.

3. Ostrom QT, Cioffi G, Gittleman H, Patil N, Waite K, Kruchko C, et al. CBTRUS Statistical Report: Primary Brain and Other Central Nervous System Tumors Diagnosed in the United States in 2012–2016. Neuro-oncology. 2019;21(Supplement_5):v1–v100.

4. Villa C, Miquel C, Mosses D, Bernier M, and Di Stefano AL. The 2016 World Health Organization classification of tumours of the central nervous system. La Presse Médicale. 2018;47(11, Part 2):e187–e200.

5. Nejat F, El Khashab M, and Rutka JT. Initial management of childhood brain tumors: neurosurgical considerations. J Child Neurol. 2008;23(10):1136–48.

6. AlRayahi J, Zapotocky M, Ramaswamy V, Hanagandi P, Branson H, Mubarak W, et al. Pediatric Brain Tumor Genetics: What Radiologists Need to Know. Radiographics: a review publication of the Radiological Society of North America, Inc. 2018;38(7):2102–22.

7. Goo HW, and Ra YS. Advanced MRI for Pediatric Brain Tumors with Emphasis on Clinical Benefits. Korean J Radiol. 2017;18(1):194–207.

8. Panigrahy A, and Bluml S. Neuroimaging of pediatric brain tumors: from basic to advanced magnetic resonance imaging (MRI). J Child Neurol. 2009;24(11):1343–65.

9. Plaza MJ, Borja MJ, Altman N, and Saigal G. Conventional and advanced MRI features of pediatric intracranial tumors: posterior fossa and suprasellar tumors. AJR American journal of roentgenology. 2013;200(5):1115–24.

10. Reardon DA, and Wen PY. Glioma in 2014: unravelling tumour heterogeneity-implications for therapy. Nature reviews Clinical oncology. 2015;12(2):69–70.

11. Wang Y, Wang Q, Haldar JP, Yeh FC, Xie M, Sun P, et al. Quantification of increased cellularity during inflammatory demyelination. Brain: a journal of neurology. 2011;134(Pt 12):3590–601.

12. Ye Z, Price RL, Liu X, Lin J, Yang Q, Sun P, et al. Diffusion Histology Imaging Detects and Classifies Glioblastoma Pathology Missed by Conventional Magnetic Resonance Imaging. bioRxiv. 2019:843367.

13. Wang Y, Sun P, Wang Q, Trinkaus K, Schmidt RE, Naismith RT, et al. Differentiation and quantification of inflammation, demyelination and axon injury or loss in multiple sclerosis. Brain: a journal of neurology. 2015;138(Pt 5):1223–38.

14. Sun P, George A, Perantie DC, Trinkaus K, Ye Z, Naismith RT, et al. Diffusion basis spectrum imaging provides insights into MS pathology. Neurology – Neuroimmunology Neuroinflammation. 2020;7(2):e655.

15. Ye Z, George A, Wu AT, Niu X, Lin J, Adusumilli G, et al. Diffusion Histology Imaging to Improve Lesion Detection and Classification in Multiple Sclerosis. medRxiv. 2019:19009126.

16. Sun P, Murphy RKJ, Gamble P, George A, Song SK, and Ray WZ. Diffusion Assessment of Cortical Changes, Induced by Traumatic Spinal Cord Injury. Brain Sci. 2017;7(2).

17. Zhan J, Lin TH, Libbey JE, Sun P, Ye ZZ, Song CY, et al. Diffusion Basis Spectrum and Diffusion Tensor Imaging Detect Hippocampal Inflammation and Dendritic Injury in a Virus-Induced Mouse Model of Epilepsy. Frontiers in neuroscience. 2018;12.

18. Vanan MI, and Eisenstat DD. Management of high-grade gliomas in the pediatric patient: Past, present, and future. Neuro-Oncology Practice. 2014;1(4):145–57.

19. Pfister S, Hartmann C, and Korshunov A. Histology and Molecular Pathology of Pediatric Brain Tumors. Journal of Child Neurology. 2009;24(11):1375–86.

20. McGirt MJ, Villavicencio AT, Bulsara KR, and Friedman AH. MRI-guided stereotactic biopsy in the diagnosis of glioma: comparison of biopsy and surgical resection specimen. Surgical neurology. 2003;59(4):277–81; discussion 81-2.

21. Apuzzo ML, Chandrasoma PT, Cohen D, Zee CS, and Zelman V. Computed imaging stereotaxy: experience and perspective related to 500 procedures applied to brain masses. Neurosurgery. 1987;20(6):930–7.

22. Air EL, Warnick RE, and McPherson CM. Management strategies after nondiagnostic results with frameless stereotactic needle biopsy: Retrospective review of 28 patients. Surgical neurology international. 2012;3(Suppl 4):S315–9.

23. Mabray MC, Barajas RF, Jr., and Cha S. Modern brain tumor imaging. Brain Tumor Res Treat. 2015;3(1):8–23.

24. Villanueva-Meyer JE, Mabray MC, and Cha S. Current Clinical Brain Tumor Imaging. Neurosurgery. 2017;81(3):397–415.

25. Kumar AJ, Leeds NE, Fuller GN, Van Tassel P, Maor MH, Sawaya RE, et al. Malignant gliomas: MR imaging spectrum of radiation therapy-and chemotherapy-induced necrosis of the brain after treatment. Radiology. 2000;217(2):377–84.

26. Price SJ, Burnet NG, Donovan T, Green HA, Peña A, Antoun NM, et al. Diffusion tensor imaging of brain tumours at 3T: a potential tool for assessing white matter tract invasion? Clin Radiol. 2003;58(6):455–62.

27. Scherer HJ. Structural Development in Gliomas. The American Journal of Cancer. 1938;34(3):333–51.

28. Kono K, Inoue Y, Nakayama K, Shakudo M, Morino M, Ohata K, et al. The role of diffusion-weighted imaging in patients with brain tumors. AJNR American journal of neuroradiology. 2001;22(6):1081–8.

29. Gajjar A, Pfister SM, Taylor MD, and Gilbertson RJ. Molecular insights into pediatric brain tumors have the potential to transform therapy. Clinical cancer research: an official journal of the American Association for Cancer Research. 2014;20(22):5630–40.

30. Pollack IF, Agnihotri S, and Broniscer A. Childhood brain tumors: current management, biological insights, and future directions. 2019;23(3):261.

31. Gauvain KM, McKinstry RC, Mukherjee P, Perry A, Neil JJ, Kaufman BA, et al. Evaluating Pediatric Brain Tumor Cellularity with Diffusion-Tensor Imaging. Am J Roentgenol. 2001;177(2):449–54.

32. Eidel O, Burth S, Neumann J-O, Kieslich PJ, Sahm F, Jungk C, et al. Tumor Infiltration in Enhancing and Non-Enhancing Parts of Glioblastoma: A Correlation with Histopathology. PloS one. 2017;12(1):e0169292.

33. Yushkevich PA, Piven J, Hazlett HC, Smith RG, Ho S, Gee JC, et al. User-guided 3D active contour segmentation of anatomical structures: Significantly improved efficiency and reliability. NeuroImage. 2006;31(3):1116–28.

34. Chawla NV, Bowyer KW, Hall LO, and Kegelmeyer WP. SMOTE: Synthetic minority over-sampling technique. J Artif Intell Res. 2002;16:321–57.

35. Haibo H, Yang B, Garcia EA, and Shutao L. 2008 IEEE International Joint Conference on Neural Networks (IEEE World Congress on Computational Intelligence). 2008:1322–8.

36. Mart, í, Abadi n, Barham P, Chen J, Chen Z, et al. Proceedings of the 12th USENIX conference on Operating Systems Design and Implementation. Savannah, GA, USA: USENIX Association; 2016:265–83.

37. Terrier L-M, Bauchet L, Rigau V, Amelot A, Zouaoui S, Filipiak I, et al. Natural course and prognosis of anaplastic gangliogliomas: a multicenter retrospective study of 43 cases from the French Brain Tumor Database. Neuro-Oncology. 2017;19(5):678–88.

38. Ruopp MD, Perkins NJ, Whitcomb BW, and Schisterman EF. Youden Index and optimal cut-point estimated from observations affected by a lower limit of detection. Biom J. 2008;50(3):419–30.

39. Carpenter J, and Bithell J. Bootstrap confidence intervals: when, which, what? A practical guide for medical statisticians. Stat Med. 2000;19(9):1141–64.

40. Pedregosa F, Varoquaux G, Gramfort A, Michel V, Thirion B, Grisel O, et al. Scikit-learn: Machine Learning in Python. J Mach Learn Res. 2011;12:2825–30.

